# The ancestral *C. elegans* cuticle suppresses *rol-1*

**DOI:** 10.1101/2020.02.07.938696

**Authors:** Luke M. Noble, Asif Miah, Taniya Kaur, Matthew V. Rockman

**Affiliations:** Institut de Biologie, École Normale Supérieure, CNRS 8197, Inserm U1024, PSL Research University, F-75005 Paris, France; Center for Genomics and Systems Biology, Department of Biology, New York University, New York, NY, 10003, USA

**Keywords:** genetic background, genetic interaction, cuticle, collagen

## Abstract

Genetic background commonly modifies the effects of mutations. We discovered that worms mutant for the canonical *rol-1* gene, identified by Brenner in 1974, do not roll in the genetic background of the wild strain CB4856. Using linkage mapping, association analysis and gene editing, we determined that N2 carries an insertion in the collagen gene *col-182* that acts as a recessive enhancer of *rol-1* rolling. From population and comparative genomics, we infer the insertion is derived in N2 and related laboratory lines, likely arising during the domestication of *Caenorhabditis elegans*, and breaking a conserved protein. The ancestral version of *col-182* also modifies the phenotypes of four other classical cuticle mutant alleles, and the effects of natural genetic variation on worm shape and locomotion. These results underscore the importance of genetic background and the serendipity of Brenner’s choice of strain.

## Introduction

Since Morgan’s first white-eyed fly, forward genetics has been one of our most powerful tools for discovering biological mechanisms. In 1974, Sydney Brenner introduced geneticists to *C. elegans*, an experimental organism with properties ideal for probing the molecular basis of development and neurobiology (Brenner 1974). Brenner began by isolating mutants with conspicuous effects under the evocative nomenclature of Dumpy, Squat, Long, Blistered, and Roller phenotypes, including a single allele of *rol-1*. This mutation causes helical twisting of the adult worm’s cuticle, which manifests most obviously as sinusoidal motion along the short axis of locomoting animals and consequent gyration on the uniform surface of an agar plate.

*rol-1*, and several other genes from Brenner’s first screen, opened the door not only to linkage mapping in *C. elegans* but also to decades of productive work on the worm cuticle. The cuticle is a complex structure, made primarily of cross-linked collagens generated anew with each larval molt (Page and Johnstone 2007). It plays an integral structural role as both barrier and morphological scaffold for muscle attachment. Epistasis analysis of collagens and collagen-modifying enzymes represents a landmark example of the power of transmission genetics to reveal molecular and developmental mechanisms (Higgins and Hirsh 1977; Cox *et al.* 1980; Kramer and Johnson 1993; McMahon *et al.* 2003).

Brenner’s original screen, and the vast majority of subsequent research in *C. elegans*, took place in the genetic context of the inbred reference strain, N2. Over the past decade, researchers have discovered that N2 evolved during its adaptation to laboratory conditions, and that wild isolates of *C. elegans* differ from the lab strain in diverse and substantive ways (Hodgkin and Doniach 1997; de Bono and Bargmann 1998; McGrath *et al.* 2009; Duveau and Félix 2012; Andersen *et al.* 2014; Sterken *et al.* 2015; Large *et al.* 2016; Gimond *et al.* 2019). Critically, the effects of mutations are often modified by genetic background. This kind of background dependence both complicates experimental analyses and underlies important genetic phenomena such as variable penetrance of Mendelian diseases in humans (Summers 1996; Scriver and Waters 1999; Dipple and McCabe 2000; Gibson and Dworkin 2004; Paaby and Rockman 2014; Paaby and Gibson 2016).

While using a *rol-1* allele as a visible marker for genetic mapping experiments, we discovered that its rolling phenotype is substantially suppressed by a wild strain background. We mapped the major-effect locus responsible for this suppression, finding that N2 carries a derived insertion in *col-182*, a collagen gene with no known mutational effects. The ancestral allele of this collagen, found in all wild isolates of *C. elegans* and highly conserved among *Caenorhabditis* species, also modifies the effects of other canonical cuticle mutants, including those with Blistered and Squat phenotypes. These results underscore the importance of genetic background and the serendipity of Brenner’s choice of strain.

## Materials and Methods

### Strains

All experiments were carried out at 20° with NGM-agarose plates and OP50-1 *E. coli* for food, unless otherwise noted.

We used the following strains: BE8: *sqt-3(sc8)* V, BE13: *sqt-1(sc13)* II, BE22: *rol-1(sc22)* II, BE44: *dpy-8(sc44)* X, BE93: *dpy-2(e8)* II, BE108: *sqt-2(e108)* II, CB61: *dpy-5(e61)* I, CB91: *rol-1(e91)* II, CB224: *dpy-11(e224)* V, CB768: *bli-2(e768)* II, CB769: *bli-1(e769)* II, CB1166: *dpy-4(e1166)* IV, CB2070: *bli-1(e935) rol-1(e91)* II, CB4856: Hawaiian wild type, COP1834: *col-182(knu732)* X, EG7993: *oxTi412 [eft-3p::TdTomato::H2B]* X, EG8951: *oxTi1015 [eft-3p::GFP::NLS + NeoR]* X, N2: laboratory wild type, QG2797: *rol-1(e91)* II; *ajIR6 [X, CB4856* > *N2]* X, QG2798: *rol-1(e91)* II; *oxTi412 [eft-3p::TdTomato::H2B] oxTi1015 [eft-3p::GFP::NLS + NeoR]* X, QG2804: *mIs12 rol-1(e91)* II, QG2952: *bli-1(e769)* II; *col-182(knu732)* X, QG2954: *dpy-11(e224)* V; *col-182(knu732)* X, QG2953: *rol-1(e91)* II; *col-182(knu732)* X, QG2955: *dpy-4(e1166)* IV; *col-182(knu732)* X, QG2956: *bli-1(e935) rol-1(e91)* II; *col-182(knu732)* X, QG2957: *rol-1(sc22)* II; *col-182(knu732)* X, QG2958: *dpy-2(e8)* II; *col-182(knu732)* X, QG2960: *dpy-5(e61)* I; *col-182(knu732)* X, QG2961: *sqt-2(e108)* II; *col-182(knu732)* X, QG2962: *bli-2(e768)* II; *col-182(knu732)* X, QG3070: *sqt-3(sc8)* V; *col-182(knu732)* X, QG3072: *sqt-1(sc13)* II; *col-182(knu732)* X, QG3074: *dpy-10(cn64)* II; *col-182(knu732)* X, QG3076: *dpy-8(sc44) col-182(knu732)* X, SP419: *unc-4(e120) rol-1(e91)* II, TN64: *dpy-10(cn64)* II, and WE5241: *ajIR6 [X, CB4856* > *N2]* X. COP1834 was generated by Knudra Biosciences (now NemaMetrix).

We verified the presence of the *rol-1(e91)* mutation in these strains by Sanger sequencing using primers RolF3: CAAATTC-GACAAAGCGACAA and RolR3: GAGCATCGTAAGGCTG-GAAA. We verified the presence of the *col-182(knu732)* mutation by Sanger sequencing using primers X12.636F: TAG-GCAAACTTGCTGCACAC and X12.636R: GAGACAGGCTG-GAAATGAGC.

### Observation of segregation distortion

As described in Kaur and Rockman (2014), we crossed wild isolate CB4856 and a strain carrying *unc-4(e120) rol-1(e91)* II in the N2 background. *F*_1_ hermaphrodites were singled and Rol nonUnc *F*_2_ adults were isolated and genotyped by Illumina GoldenGate Assay. Genotyping was performed by the DNA Sequencing and Genomics Core Facility of the University of Utah. Allele frequencies are in File S2.

### Complementation crosses

We used visible markers to generate animals that are homozygous *rol-1* on chromosome II and heterozygous on the X chromosome, with one X chromosome from N2 and the second from the strain of interest (SOI). If the SOI carries the dominant suppressor of *rol-1*, then these animals are expected to show suppressed rolling behavior.

Specifically, we crossed *mIs12[GFP] rol-1* II males to SOI hermaphrodites to generate *mIs12 rol-1 / + +* II; SOI X *F*_1_ males. These we crossed to *unc-4 rol-1* II hermaphrodites. For each SOI, we tested six GFP-positive nonUnc hermaphrodite progeny of this cross for rolling behavior in the second day of adulthood. These animals are mostly *unc-4 + rol-1 / + mIs12 rol-1* II; SOI/N2 X. A small fraction of the phenotyped animals could be *rol-1* heterozygotes due to rare recombination events between *mIs12* and *rol-1* in the *F*_1_ males. In addition, each tested animal is heterozygous (N2/SOI) for a random fraction (expectation ½) of the autosomes except for chromosome II.

We also tested 8 wild isolates by this approach (CB4856, PB306, EG4347, QX1211, JU319 [CeNDR isotype JU311], JU1088, PX179, and JU400 [CeNDR isotype JU394]), along with N2 and LJS1, which are laboratory-adapted strains.

### Fine mapping with recombinants

To fine-map the suppressor, we crossed *rol-1(e91)* II; CB4856 X and *rol-1(e91)* II; *oxTIi1015 oxTi412* X. The latter strain carries integrated single-copy transgenes on the X expressing fluorescent proteins, tdTomato::H2B from X:11.049 Mb and GFP::NLS from X:13.480 Mb [Wormbuilder; Frøkjær-Jensen *et al.* (2014)]. Both strains carry N2-derived autosomes. We homozygosed X chromosomes that were recombinant between the transgenes, scored their rolling behavior, and genotyped them at SNP markers in the mapping interval. Recombinant strain AM_GNR9 placed the suppressor to the right of SNP WBVar00083599 at X:12,579,121, and recombinant AM_GNR13 placed it to the left of SNP WBVar1602269 at X:12,662,633.

Genotyping primers used for mapping were: WBVar01981458 indel, TGGGTAAACATCGGCTCCAT and TGTTCTGCACGGGAAAAGAT; WBVar00083599 SNP, CGACATCCAAAGTTTTTGAGACT and GAGAAAGTGT-TATGGGCATGG; WBVar01602269 SNP, CGTGTGTTTC-CGTTGTGAAT and TTCAGTGTTCATCGCAATCTG.

### Association mapping

Variant data for 330 *C. elegans* isotypes were downloaded as the 20180527 CeNDR release soft-filtered vcf. We tested for a match between variants in the recombination-mapping interval (X:12,579,122 - X:12,662,632) and *rol-1* suppression phenotype in the panel of N2, CB4856, LSJ1, and the seven other tested wild isolates.

### Gene model

Sequence and annotation data for the *Caenorhabditis* genus were downloaded from the *Caenorhabditis* Genomes Project. RNAseq data from young adults of *C. elegans* wild isolates were downloaded from the NCBI SRA: CB4856 PRJNA437313 (Zamanian *et al.* 2018), AB1 and ED3040 PRJNA288824 (Vu *et al.* 2015). Reads were mapped to a 100 Kb region of the N2 WS220 genome centered on *col-182* with bwa mem version 0.7.17 (Li and Durbin 2010), assembled with Trinity version 2.3.2 in genome-guided mode (Grabherr *et al.* 2011), and homologous transcripts were extracted by blast version 2.9 (Altschul *et al.* 1990) against *col-182* orthologs across the genus. Coding and protein sequences were aligned with mafft version 7.3.10 in L-INS-i mode (Katoh *et al.* 2005), and homology and gene structures were plotted using R packages *ape* (Paradis and Schliep 2019), *ggtree* (Yu *et al.* 2017), *Biostrings* (Pagès *et al.* 2019), and *ggbio* (Yin *et al.* 2012). Species with only computationally predicted annotations (*C. kamaaina, C. becei, C. panamensis*) and two species (*C. quiockensis* and *C. sulstoni*) with extremely long predicted ortholog sequences potentially deriving from annotation errors were excluded. Collagen triplet stability scores shown in Figure 2B were obtained from (Persikov *et al.* 2005), and ignore higher-order interactions. Gene models and coding sequence alignments are in Files S10,S11.

**Figure 1.**
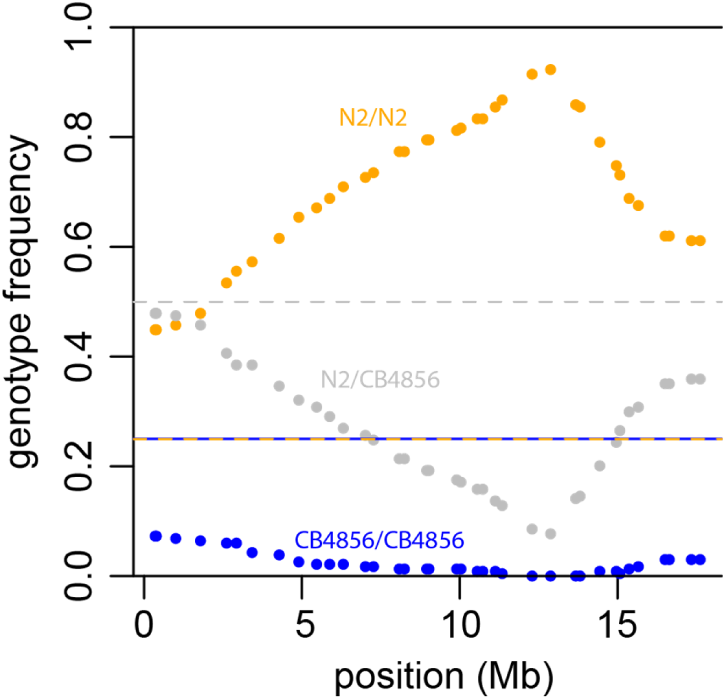
Allele frequencies along the X chromosome in Rol nonUnc *F*_2_s from a cross of CB4856 and N2-background *unc-4(e120) rol-1(e91)* II. Dashed lines show the Mendelian expectations for heterozygotes (½) and homozygotes (¼ each).

**Figure 2.**
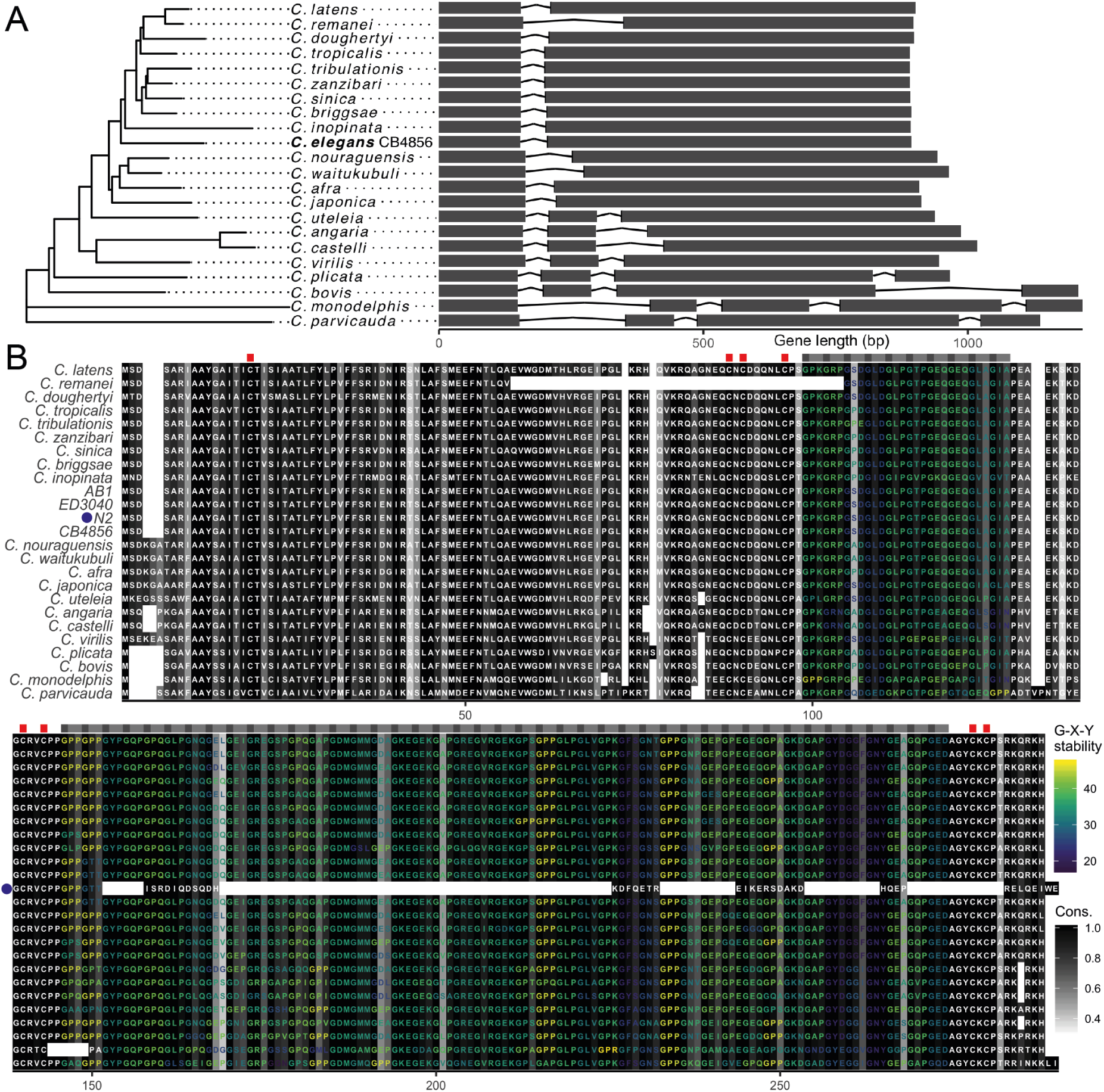
A. *col-182* gene tree and model for *C. elegans* CB4856 and other *Caenorhabditis* species. The x-axis shows distance from the start codon for each genome. **B**. Protein alignment, including predicted sequences for *C. elegans* wild isolates CB4856, AB1 and ED3040 assembled from young adult RNAseq data, and the frame-shifted N2 translation (marked with a blue dot). Grey-scale shading represents site conservation (% identity), and labels are colored by predicted collagen triplet stability (melting temperature) for runs of >1 G-X-Y repeats conserved across all sequences other than N2. The positions of conserved triplets are indicated above the alignment by grey boxes, and the positions of conserved cysteine residues potentially involved in inter-strand disulphide bridges are shown as red boxes.

### Epistasis analysis of visible mutants

For each tested mutation, we grew the mutant line and *col-182(knu732)* double mutant in parallel at low population densities for several generations and performed synchronous egg lays to generate animals for phenotyping. For most strains we 2 Noble *et al.* observed these synchronized animals three days later as young adults. For *sqt-2(sc108)*, because the heterozygous phenotype (Rol) is different from the homozygous phenotype (Sqt), we also scored *sqt-2/+* animals. For *rol-1 bli-1* and *rol-1 bli-1; col-182* animals, we followed 200 of each genotype through the third day of adulthood. Videos of *rol-1* worms (CB91, QG2979, QG2957, BE22, QG2953) were taken on day four of adulthood. Ten worms were picked to fresh seeded plates and imaged for 8 minutes at 12 frames per minute. Videos of *sqt-3* worms (BE8, QG3070) were taken on day two of adulthood. Twenty worms were picked to fresh seeded plates and imaged for five minutes at 12 frames per minute. Video samples are in Files S3-S9.

### Quantitative locomotion analysis

Conditions and worm tracking have been described previously (Noble *et al.* 2017; Mallard *et al.* 2019). Lines (N2, COP1834, BE8, BE22, QG2957, QG3070) were each split to duplicate lineages, bleached, and grown under common conditions on HT115 bacteria in 90mm plates at 20° for two generations before assay, with each generation starting from around 500 L1 larvae that had been starvation synchronised in M9 buffer for 18 hours. Young adults were imaged in random order, on food, during day three post-L1, and again on two further generations (treated as above) for a total of six replicate plates per genotype. Worms were tracked for 8 minutes using the Multi-Worm Tracker (Swierczek *et al.* 2011), the final 4 of which were analysed after subsampling to 4 Hz, and 11-point skeletons and outlines from Choreography were parsed to generate summary track statistics based on size and movement. More precisely, we used Choreography-defined measurements of length, width, area, speed, acceleration, angular momentum (turning rate), mean body curvature, kink (the maximum ratio of angles between head/tail and body), and the length of continuous runs of Forward, Backward or Still motion (“bias”). Raw data are in File S12.

For each worm track we took the median and variance of each metric, as a whole and split by bias state. Exploratory behaviour was quantified as the area and circularity (4*π area*/*perimeter*^2^) of the track convex hull (Pebesma and Bivand 2005), averaged first across 30 second intervals for each of the longest 100 tracks from each plate, then across tracks. Traits were log transformed where strongly non-normal (an improvement in Shapiro-Wilk −log10 p-value > 6), and effects of assay block, defined by lineage and assay day, were removed by linear regression. Processed data are in File S13.

We show univariate and multivariate (classical multidimensional scaling on the mean centered and scaled Euclidean distance matrix of plate means, base R *cmdscale*) analysis. In Figure 3A-B we used multivariate analysis of a subset of seven traits selected using sparse linear discriminant analysis (Clemmensen *et al.* 2011). From 25 traits (repeatability > 0.5, thinned to reduce maximum colinearity to *r*^2^ < 0.5), we selected the five metrics most associated with suppression of *rol-1(sc22)* or *sqt-3(sc8)*, separately, using plate means. Traits retained for *sqt-3* (BE8 vs. QG3070, COP1834, N2), ordered by absolute loading on the discriminant function, were log transformed curvature (S), width (F), kink, width (S) variance, and acceleration (F). Traits retained for *rol-1* (BE22 vs. QG2957, COP1834, N2) were log transformed circularity, velocity variance, width (F), kink (F), and run length (F) variance. R code for this analysis is in File S14.

**Figure 3.**
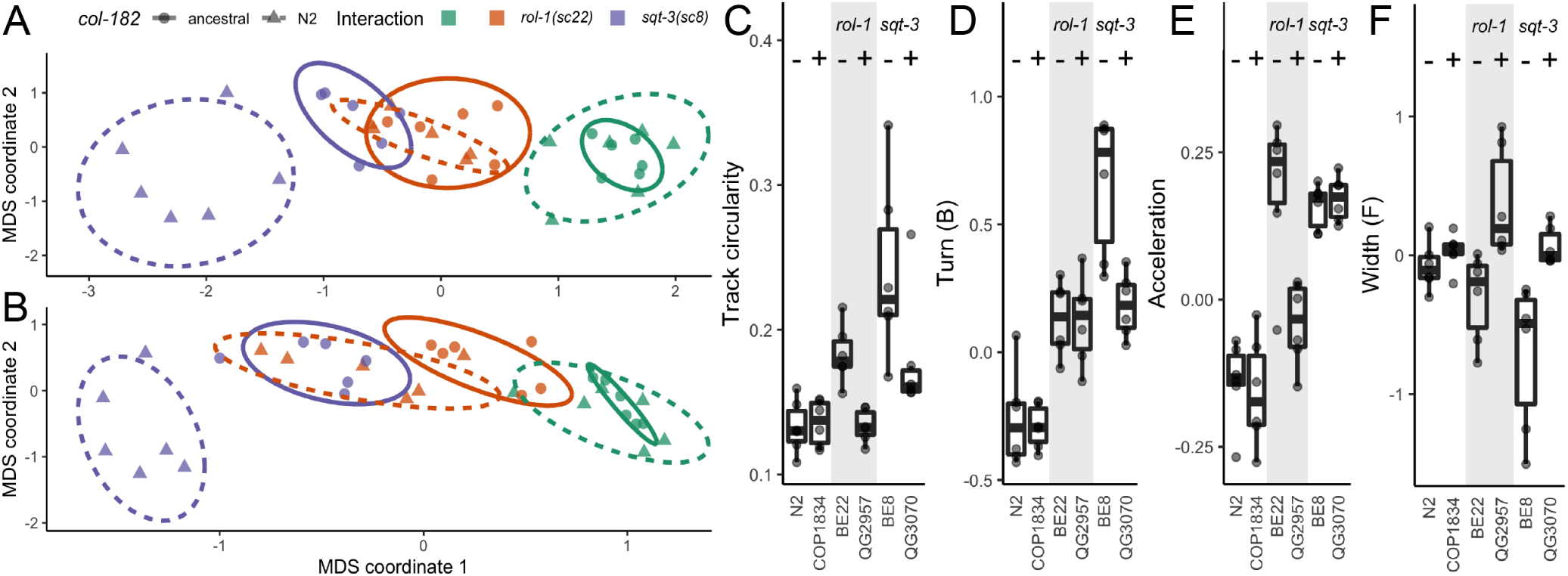
Ancestral *col-182* suppresses *rol-1* and *sqt-3* alleles. **A-B**. Multidimensional scaling of locomotion and size traits: an unbiased set of 19 traits selected only on repeatability across replicate plates (**A**) and a set of seven traits selected by sparse discriminant analysis that maximise multivariate suppression of *sqt-3* or *rol-1* by ancestral *col-182* (**B**). Each point is the grand mean of tracks from around 500 young adult worms per replicate plate over three consecutive generations for each genotype, assayed for N2 (green triangles) and COP1834 (ancestral *col-182* in the N2 background; green circles), BE22 (N2 *col-182; rol-1(sc22)*; orange triangles) and QG2957 (ancestral *col-182; rol-1(sc22)*; orange circles), and BE8 (N2 *col-182; sqt-3(sc8)*; purple triangles) and QG3070 (ancestral *col-182; sqt-3(sc8)*; purple circles). **C-F**. Univariate comparisons show variable effects across backgrounds. *col-182* genotype is indicated with – (N2 insertion) and + (ancestral) symbols. Complete suppression of track circularity is seen for *rol-1(sc22)* (**C**), and of worm width in the forward state for *sqt-3(sc8)* (**F**), however partial (or no) suppression is the most common outcome. Raw and processed data in Files S12,S13, code in File S14.

### Epistasis analysis in the CeMEE

To test for potential modifying effects of *col-182* on natural variation segregating in the *C. elegans* multiparent experimental evolution (CeMEE) panel we genotyped the N2 indel from existing sequence data (Noble *et al.* 2019) using bcftools (Li 2011) after indel realignment (DePristo *et al.* 2011), obtaining calls for 365 recombinant inbred lines (RILs; from populations A6140, CA[1-3]50 and GA[1,2,4]50) for which locomotion has been measured on NGM (Mallard *et al.* 2019). Two lines were excluded as multivariate outliers based on Mahalanobis distance. Genotypes were marker set 1 from Noble *et al.* (2019). The N2 *col-182* allele is at a frequency of 16.5% in these lines, providing sufficient power to detect pairwise interactions conditional on joint allele frequency. In total, 167,187 diallelic SNPs where all four genotype classes were present at a minimum frequency of 10, excluding any uncertain imputations, were tested.

To test for *col-182*-by-genotype interactions we fit nested bivariate linear models for three pairs of partially correlated traits: length and width (forward state, log transformed), the *rol-1* and *sqt-3* suppression discriminant functions that are linear combinations of five traits (see Quantitative locomotion analysis), and the single traits with the highest loading in each discriminant function, curvature and track circularity (log transformed). Trait values were best linear unbiased predictions (BLUPs) extracted from linear mixed effects models (R package *lme*4) fit to replicate observations, with fixed effects of population replicate. Significance testing followed the univariate approach in Noble *et al.* (2017). In brief, we first tested for genetic effects by likelihood ratio (Pillai’s trace statistic) for a full mode, with additive and interaction effects of *col-182* genotype and focal marker genotype, against a null model (intercept only). Genome-wide significance was declared against a null distribution of >1000 test statistics generated by permuting lines within populations and retaining the minimum observed *p*-value. We used a relative permissive false discovery rate (FDR) of 0.2. Quantile-quantile plots showed statistics were well calibrated for length/width and the Rol/Sqt discriminant functions at *p* > 10^−3^, but strongly deflated for curvature/circularity. Inflation was evident for length/width at *p* < 10^−3^, independent of linkage disequilibrium, consistent with additional polygenic interactions. For loci with significant genetic effects, interaction significance was then assessed at a nominal threshold of *p* < 0.05 by parametric bootstrap against the additive model (Bůžková *et al.* 2011), resampling responses jointly among lines 5000 times. Genotype and phenotype data, and R code for this analysis are in Files S15,S16.

### Gene expression analysis

We extracted data for 199 recombinant inbred advanced intercross lines (RIAILs) from Rockman *et al.* (2010), after excluding data from lines with annotation issues (Zych *et al.* 2017). We performed structured nonparametric trait mapping as in Rockman and Kruglyak (2009) for abundance of 15,617 transcripts whose genes are more than a megabase from *col-182* (to exclude local linkages for genes near *col-182* that have their own *cis*-acting variants). We retained traits with genome- and experiment-wide significant linkage peaks (LOD>4.3, 5% FDR) within 1 RIAIL-effective cM of *col-182* (approximately 400 Kb). The nine significantly linked transcripts were tested for functional enrichment using the WormBase Enrichment Analysis Suite (Angeles-Albores *et al.* 2018).

### Data Availability

All quantitative data and code to reproduce our analyses and main figures are available from FigShare (XXX) and github.com/lukemn/cuticle. Strains are available upon request. File S1 details all supplemental files, File S2 contains genotyping data from Figure 1, Files S3-S9 contain short videos of mutant and suppressed adult hermaphrodites on plates, File S10 contains gene models from Figure 2A, File S11 contains coding sequence alignments for Figure 2B, Files S12 and S13 contain raw and processed Multi-Worm tracker data for Figure 3 with associated R code in File S14. File S15 contains genotypes and phenotypes used for detecting genetic interactions between *col-182* and SNPs in the CeMEE (Figure 4), with associated R code in File 16.

**Figure 4.**
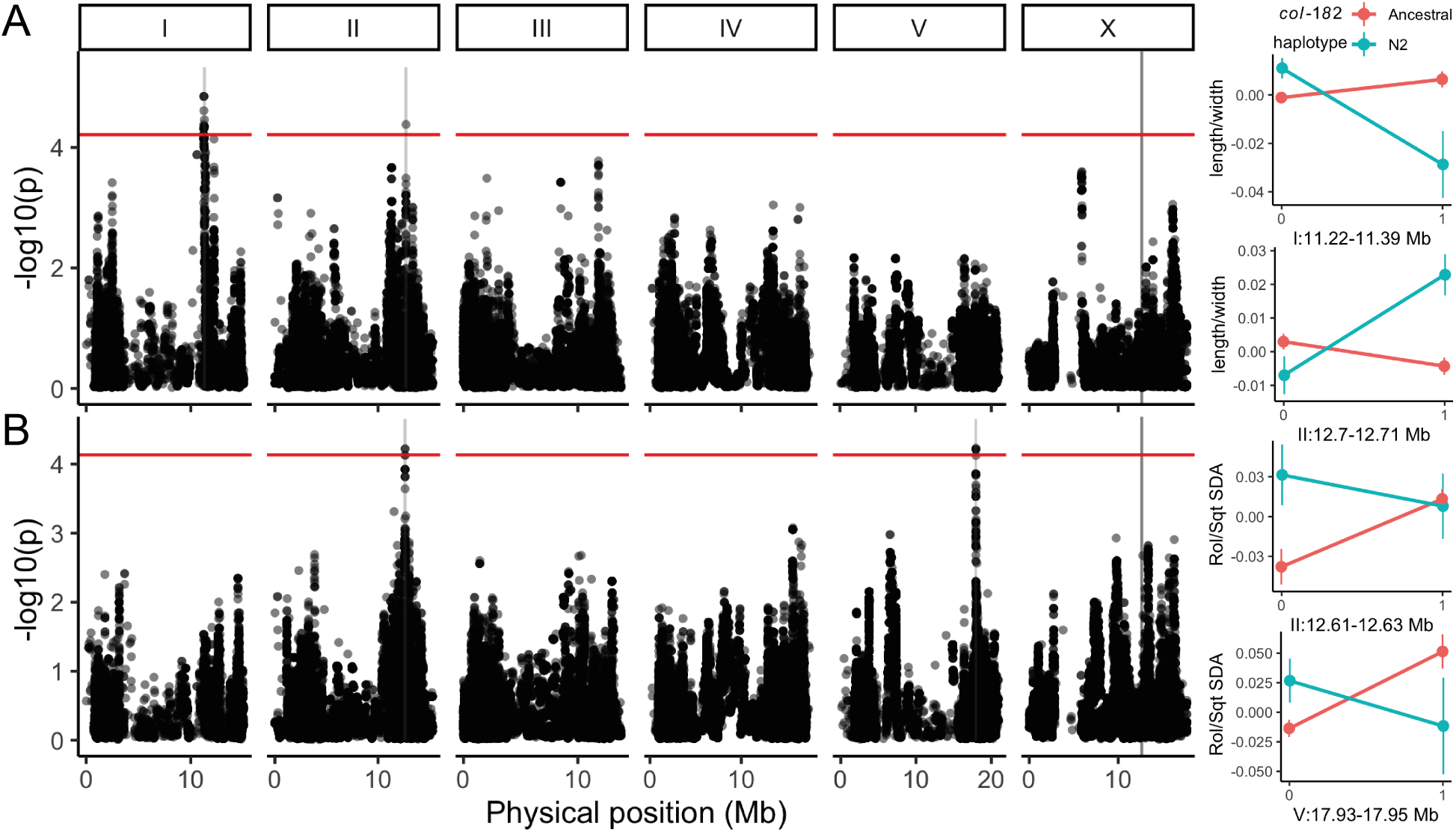
*col-182* modifies the effects of natural genetic variation on worm movement and shape. Genome-wide statistics are shown for two bivariate response models of *col-182* indel × SNP genotype, worm length/width (**A**) and Rol/Sqt sparse discriminant functions (**B**), using diallelic SNPs segregating in 363 recombinant inbred lines of the *C. elegans* multiparent experimental evolution (CeMEE) panel. Statistics are from a likelihood ratio test for a full additive and interaction linear model against a null model of no genetic effects. Permutation thresholds for genome-wide significance (FDR = 0.2) are shown in red, pink shaded regions show 1.5 LOD drop QTL intervals (expanded to a minimum of 300 Kb for visibility), and the location of *col-182* on the X chromosome is indicated by a grey line. Effect plots at right show genotype class means and standard errors for QTL with significant interactions (parametric bootstrap against additive models, *α* = 0.05). Trait values shown are the first principal component for length/width, and the Sqt discriminant function, which explains most of the interaction in the Rol/Sqt bivariate model. Reference-based genotype is on the x-axis. Genotype and phenotype data in File S15, code in File S16.

## Results

### CB4856 carries a dominant suppressor of rol-1

During a study of recombination patterns on *C. elegans* chromosome II (Kaur and Rockman 2014), we crossed *unc-4(e120) rol-1(e91)* II worms (N2 background) with wild isolate strain CB4856, isolated in Hawaii in 1972 (Hodgkin and Doniach 1997). After allowing *F*_1_ hermaphrodites to reproduce by selfing, we isolated Rol nonUnc recombinants, which are homozygous for the N2 *rol-1* allele but carry at least one CB4856 allele at *unc-4* and should show Mendelian segregation of other chromosomes (*pace* the *peel-1 zeel-1* incompatibility on chromosome I (Seidel *et al.* 2008)). We noticed strong segregation distortion on the X chromosome among Rol nonUnc recombinants, which we had genotyped at 37 SNP markers (Figure 1, File S2). Distortion favored the N2 background, with a peak around 13 Mb where zero of 234 worms were homozygous for the CB4856 genotype (expectation ¼ = 58.5). At the peak, 216 (92%) worms were N2 homozygotes and the remaining 18 (8%) were heterozygotes.

We hypothesized that the CB4856 X chromosome carries a dominant suppressor of *rol-1*. We crossed the *rol-1(e91)* allele into an X-chromosome substitution strain, which carries the CB4856 X chromosome but is otherwise N2, and confirmed that the resulting *rol-1(e91)* II CB4856 X strain is strongly, but incompletely, suppressed for rolling. The worms retain a slight helical twist and can be distinguished from wildtype N2 and CB4856, but the dramatic rolling and circling behaviors of *rol-1(e91)* are absent (Videos: CB91 vs. QG2797 in Files S3,S4). Suppression of *rol-1* is also observed in X chromosome heterozygotes, consistent with a dominant suppressor on the CB4856 X and explaining the pattern of segregation distortion we observed in the *unc-4 rol-1* crosses.

Suppression often reveals interactions between genes in physical association or in specific developmental pathways, but can also reflect altered transcription, splicing, or translation of a mutant gene. These informational suppressors can be allele specific, masking only particular kinds of missense or nonsense or splice site variants, for example (Hodgkin 2005). There are two *rol-1* alleles with known effects on phenotype at present, and we found that the second, *sc22*, is also suppressed by the CB4856 X chromosome. Although the *sc22* molecular lesion is unknown, *e91* and *sc22* mutants show distinct phenotypic profiles, including temperature sensitivity (Cox *et al.* 1980), and we concluded that allele-specific interaction was thus unlikely.

### The suppressor maps to an indel polymorphism in col-182

We performed complementation testing with a panel of N2-CB4856 recombinant inbred advanced intercross lines (Rockman and Kruglyak 2009) with breakpoints near the peak allele frequency distortion, taking advantage of the dominant mode of action of the CB4856 allele. Using this approach, we tested 5 RIAILs, scoring based on Rolling, with boundaries defined by RIAIL QX43, which carries the suppressor, and RIAIL QX126, which does not. These strains place the suppressor to the right of SNP WBVar00083496 at X:12,364,484 and to the left of indel WBVar01981458 at X:12,699,819 (WS272 coordinates).

Next we used integrated single-copy fluorescent marker transgenes (Frøkjær-Jensen *et al.* 2014) to select for N2/CB4856 recombinants in the interval, in N2 autosomal genetic backgrounds. These data localized the causative locus to the interval between 12.579 and 12.662 Mb.

Complementation testing with seven additional wild isolates found that all exhibit suppression of *rol-l*. Laboratory strain LSJ1, which shares a laboratory ancestor with N2 but was cultured separately since 1963, does not exhibit suppression. Among all variants segregating in these strains in the mapped interval, only one exhibits perfect cosegregation with *rol-1* suppression: an 8-base pair (bp) deletion in the gene *col-182* (WBVar01928355). Like *rol-1, col-182* is one of the 181 collagen genes in the *C. elegans* genome (Teuscher *et al.* 2019).

### N2 carries a derived insertion mutation in col-182

Among 330 genome-sequenced *C. elegans* isotypes from around the world (CeNDR freeze 20180527), the 8-bp deletion is present in every strain except for the lab strains N2, LSJ1, and ECA252. These three strains are all derived from a single isolate of *C. elegans* sampled by L. N. Staniland in 1951 (McGrath *et al.* 2009; Weber *et al.* 2010; Sterken *et al.* 2015; Cook *et al.* 2017). The deletion state is also found in orthologs of all other examined *Caenorhabditis* species (Figure 2). This strongly suggests the reference allele is a derived insertion, one that arose either in the wild, and happened to be sampled by Staniland in 1951, or in the lab sometime before 1963 [likely before 1958 (Gimond *et al.* 2019)].

*C. elegans* collagens contain two blocks of Gly-X-Y repeats, flanked and separated by short cysteine-containing domains involved in interchain disulphide bonds (Page and Johnstone 4 Noble *et al.* 2007). Until recently, *col-182* was annotated as a three-exon, two-intron gene encoding a near-canonical collagen protein, missing only the middle cysteine domain (Teuscher *et al.* 2019). However, new RNAseq data suggests that the earlier gene model was erroneous, and that the N2 transcript has only one intron (Worm-Base WS274). With this RNAseq-supported gene model, the 8-bp insertion causes a frameshift that eliminates the entire second Gly-X-Y domain and final cysteine domain (Figure 2B). As a consequence, *col-182* has been reclassified as a pseudogene. The ancestral version of the gene, preserved in CB4856 and all other wild isolates, encodes a perfect canonical cuticular collagen. The C-terminal Gly-X-Y domain includes 43 consecutive Gly-X-Ys, tying it with four other genes for the longest such collagenous stretch among *C. elegans* collagens [with *col-72, col-75, col-104*, and *col-113*, none of which have known mutant phenotypes; (Teuscher *et al.* 2019)].

To test whether *col-182* is the *rol-1* suppressor and whether the ancestral allele functions via the predicted isoform, we commissioned a strain that carries the ancestral splicing and reading frame in an otherwise N2 background. CRISPR-Cas9 conversion removed the 8-bp insertion and altered 16 additional base pairs, each a synonymous third codon position change in the predicted ancestral reading frame. We then confirmed that this ancestral allele *col-182(knu732)* suppressed *rol-1*, both the *e91* and *sc22* alleles (videos: CB91 vs. QG2953 and BE22 vs. QG2957, Files S3,S5,S6,S7). From tracking of young adult hermaphrodite worms we derived an array of statistics describing locomotion, size, and shape (Quantitative locomotion analysis). Reduction of this data by multidimensional scaling into two orthogonal axes provided quantitative confirmation that the ancestral *col-182(knu732)* allele strongly, but incompletely, suppresses *rol-1* (Figure 3A). By univariate analysis, double mutants are indistinguishable from non-Rol genotypes for a single metric that captures circling locomotion, but weak to no suppression is the predominant outcome across the correlated set of single traits (Figure 3B).

### col-182 interacts with other collagen mutants

*rol-1* is part of a complex network of collagens and collagen-modifying enzymes that interact genetically, often in quite complex ways (Cox *et al.* 1980; Kusch and Edgar 1986). We therefore expected that the ancestral *col-182* might modify *rol-1*’s epistatic relationships with other genes, and might itself interact with other cuticle-specifying genes.

Blistered mutants, described along with *rol-1* in Brenner’s original screen, develop fluid-filled blisters along the adult cuticle (Brenner 1974). In the N2 background, *rol-1* suppresses the blister phenotype of *bli-1* mutants (Cox *et al.* 1980). We found that *rol-1 bli-1*; *col-182* triple mutants show the expected suppression of rolling but are also unblistered, indicating that *col-182* suppresses *rol-1*’s Rol phenotype but not its suppression of *bli-1*. Moreover, *col-182* modified *bli-1*’s phenotype by itself; blisters were largely suppressed in *bli-1; col-182* double mutant hermaphrodites. Blisters were fewer (23% vs. 92% in young adults, n=40 and 36), and were spatially restricted to the head region. Conversely, *col-182* enhances the blister phenotype of *bli-2* worms; *bli-2* young adults had discrete blisters on their bodies or heads (38/39), but a third of *bli-2; col-182* worms had single blisters that spanned the full length of the animal (9/30). The majority of *bli-2* animals (24/40) had blisters restricted to their heads, versus 7/30 for the *bli-2; col-182* worms. Like *col-182* and *rol-1, bli-1* and *bli-2* encode collagens.

Cox *et al.* (1980) identified alleles in several genes that give rise to left-handed rollers, like *rol-1(e91)*. Although we observed no gross phenotypic effect of *col-182* on *dpy-8(sc44), dpy-10(cn64)*, or *sqt-1(sc13)*, we observed partial suppression of rolling in *sqt-3(sc8)*, which we quantified by worm tracking (Figure 3, and see videos: BE8 vs. QG3070, Files S8,S9). Suppression was qualitatively and quantitatively distinct to that of *rol-1*; near complete for the single measure of worm width, but again highly variable and generally weak for other measures of locomotion and morphology (Figure 3B).

*sqt-2(sc108)* exhibits right-handed rolling as a heterozygote in the N2 background, and did so as well in the *col-182(knu732)* background. However, *sqt-2* heterozygotes showed slowed development in the N2 *col-182* background, while the ancestral allele suppressed the developmental delays. Finally, alleles of several additional genes involved in cuticle development – collagens *dpy-2(e8), dpy-4(e1166)* and *dpy-5(e61)*, and thioredoxin *dpy-11(e224)* – showed no gross phenotypic modification in the *col-182(knu732)* background.

### col-182 modifies effects of natural variation on worm shape and locomotion

Collagens are known to influence body size (Brenner 1974; Fernando *et al.* 2011; Madaan *et al.* 2018), and our locomotion analysis identified specific axes of worm size, posture and locomotion modified by *col-182* in two genetic backgrounds. We next sought to test more broadly for interactions between *col-182* and natural genetic variation for these traits in the *C. elegans* Multiparent Experimental Evolution (CeMEE) panel, a collection of recombinant inbred lines derived from the pooled standing genetic diversity of 14 wild isolates and two N2-related strains (Teotónio *et al.* 2012; Noble *et al.* 2017, 2019).

We genotyped the N2 insertion in RILs sampled from an ancestral laboratory-adapted population, A6140, and from six populations derived from A6140 that evolved under varying mating system and environment (Noble *et al.* 2017). Using Multi-Worm Tracker data for 363 lines, we fit bivariate linear models for three sets of correlated traits (length and width, body curvature and track circularity, and the Rol/Sqt discriminant functions from Figure 3B) to test for interaction effects. Univariate tests showed that *col-182* genotype had no effect on the means of these population-centred traits (0.37 < *p* < 0.76 by likelihood ratio test).

We detected four loci with significant genetic effects at a per-model false discovery rate of 20% (Figure 4). Two QTL were detected for length/width with clear genetic interactions (*p* < 0.001 by bootstrap against the additive model): the first, on chromosome I, fell within the central recombination rate domain (1.5 LOD drop interval around 170 Kb); the second, on chromosome II, was contained by a single very large N2 protein coding gene, *tbc-17*, with several missense variants, a splice-donor change, and heterozygous SNP calls suggestive of copy number variation segregating in the CeMEE founder haplotypes (Cook *et al.* 2017). *tbc-17* encodes a highly conserved predicted Rab family GTPase activator which, based on homology, may be involved in intracellular trafficking, a process critically important for collagen secretion from hypodermal cells (Roberts *et al.* 2003; Ackema *et al.* 2013). Two QTL were detected for the Rol/Sqt discriminant functions: one on chromosome II (interaction bootstrap *p* < 0.001; 18 Kb interval) contained nine N2 annotated protein-coding genes of unknown function, mostly of the nematode-specific peptide group E family; the second, on 6 Noble *et al.* chromosome V (interaction *p* < 0.02; 25 Kb interval), was a specific interaction with MY16 haplotypes, spanning predicted ubiquitin protease *usp-50* partially, and *dpy-21* fully, along with eight non-coding RNAs. *dpy-21* is a non-essential, non-condensin subunit of the dosage compensation (DC) complex (Meyer and Casson 1986), with additional DC-independent roles in gene regulation (Webster *et al.* 2013), and loss-of-function mutants show an enrichment in dysregulation of genes involved in the cuticle (Kramer *et al.* 2015).

### col-182 does not have systemic effects on gene expression

Several of the alleles that arose during *C. elegans* domestication have large and systemic effects on *C. elegans* biology. These include *npr-1* (de Bono and Bargmann 1998; McGrath *et al.* 2009; Andersen *et al.* 2014; Zhao *et al.* 2018), *nath-10* (Duveau and Félix 2012), *nurf-1* (Large *et al.* 2016), and *Y17G9B.8* (Rockman *et al.* 2010; Burga *et al.* 2018). We therefore investigated whether *col-182* is linked to systemic effects on gene expression in adult hermaphrodites, using a published dataset of gene expression in 199 N2/CB4856 recombinant inbred advanced intercross lines [RIAILs; (Rockman *et al.* 2010)]. The RIAILs provide much higher genotypic replication than a typical pairwise contrast of strains, as each *col-182* allele is homozygous in approximately half the RIAILs, but effects that map to *col-182* may be due to nearby variants in other genes. As expected, *col-182* abundance shows strong linkage to its own location. Nine other genes show significant linkage to the *col-182* region Table 1, including two glutathione S-transferases and two cytochrome P450 enzymes. However, none are known to be involved in cuticle development and collectively they show no enrichment for any particular tissue. Thus the *col-182* mutation in N2 appears to have limited effects on gene expression in young adult hermaphrodites, at least under ordinary laboratory conditions.

**Table 1.** Genes with expression linked to *col-182* in N2/CB4856 RIAILs (Rockman *et al.* 2010). Probe: Agilent microarray probe, ID: WormBase systematic identifier, Gene: common name (if any), Pos./Chr.: physical position of the transcript, QTL: genetic position of the quantitative trait locus in the RIAILs in cM (within 1 cM of *col-182*), LOD: logarithm of the odds of association between gene expression and the QTL peak.

| Probe | ID | Gene | Pos. | Chr. | QTL | LOD |
| --- | --- | --- | --- | --- | --- | --- |
| A_12_P107311 | R05F9.5 | *gst-9* | 4,893,372 | 2 | 134.26 | 8.78 |
| A_12_P120030 | B0464.1 | *dars-1* | 9,489,112 | 3 | 133.00 | 4.83 |
| A_12_P116084 | Y40B10A.2 | *comt-3* | 2,061,050 | 5 | 133.00 | 8.62 |
| A_12_P111642 | C45H4.17 | *cyp-33C2* | 2,186,492 | 5 | 134.26 | 5.34 |
| A_12_P104828 | C02A12.1 | *gst-33* | 3,467,564 | 5 | 133.00 | 8.50 |
| A_12_P115936 | F08F3.7 | *cyp-14A5* | 5,421,110 | 5 | 133.78 | 8.56 |
| A_12_P108481 | T04H1.9 | *tbb-6* | 12,263,208 | 5 | 134.26 | 6.42 |
| A_12_P111217 | C05C9.3 |  | 11,143,372 | 6 | 133.00 | 4.95 |
| A_12_P105133 | C35C5.6 | *trpp-9* | 11,569,206 | 6 | 133.78 | 8.13 |

## Discussion

The structural complexity of the nematode cuticle is reflected in its developmental and genetic regulation, and its environmental (temperature) and genetic sensitivity. Around 4% of the worm genome is dedicated to expressing, processing, and assembling the collagens, cuticulins, glycoproteins and other components of the multilayered extracellular matrix (Teuscher *et al.* 2019). Yet, of 173 predicted cuticular collagen genes, phenotypes from extensive mutagenesis screens have been detected for just 21 (Page and Johnstone 2007). To this number we can now add *col-182*, though we have also shown that even this select list might well have been shorter had not Brenner adopted N2 as the *C. elegans* reference genetic background.

In the absence of molecular and structural data, the precise role of *col-182* in the worm cuticle and its mode of interaction with other collagens remains obscure. The derived N2 insertion represents an evolved enhancer of *rol-1* rolling, and the ancestral *col-182* modifies to a variable extent the phenotypes from other classical mutant alleles of *bli-1, bli-2, sqt-2* and *sqt-3*, but not obviously those of *dpy-2, −4, −5, −8, −10* and *-11*, or *sqt-1*.

The expression of cuticular genes during worm development offers no clear insight. Of the tested genes, only *rol-1*, for which suppression by ancestral *col-182* was strongest, shows strong stage specificity, being around 30-fold enriched in L4 (i.e., when the adult cuticle is manufactured; Figure 5). But *bli-1, bli-2* and *rol-1* show generally similar patterns and levels of transcriptional activity over the life-cycle, with very low expression in embryonic and early larval stages. The Blistered phenotype is thought to be due to defects in struts linking basal and cortical layers of the adult cuticle, and of six Blistered mutants, three are enzymes rather than structural components.

**Figure 5.**
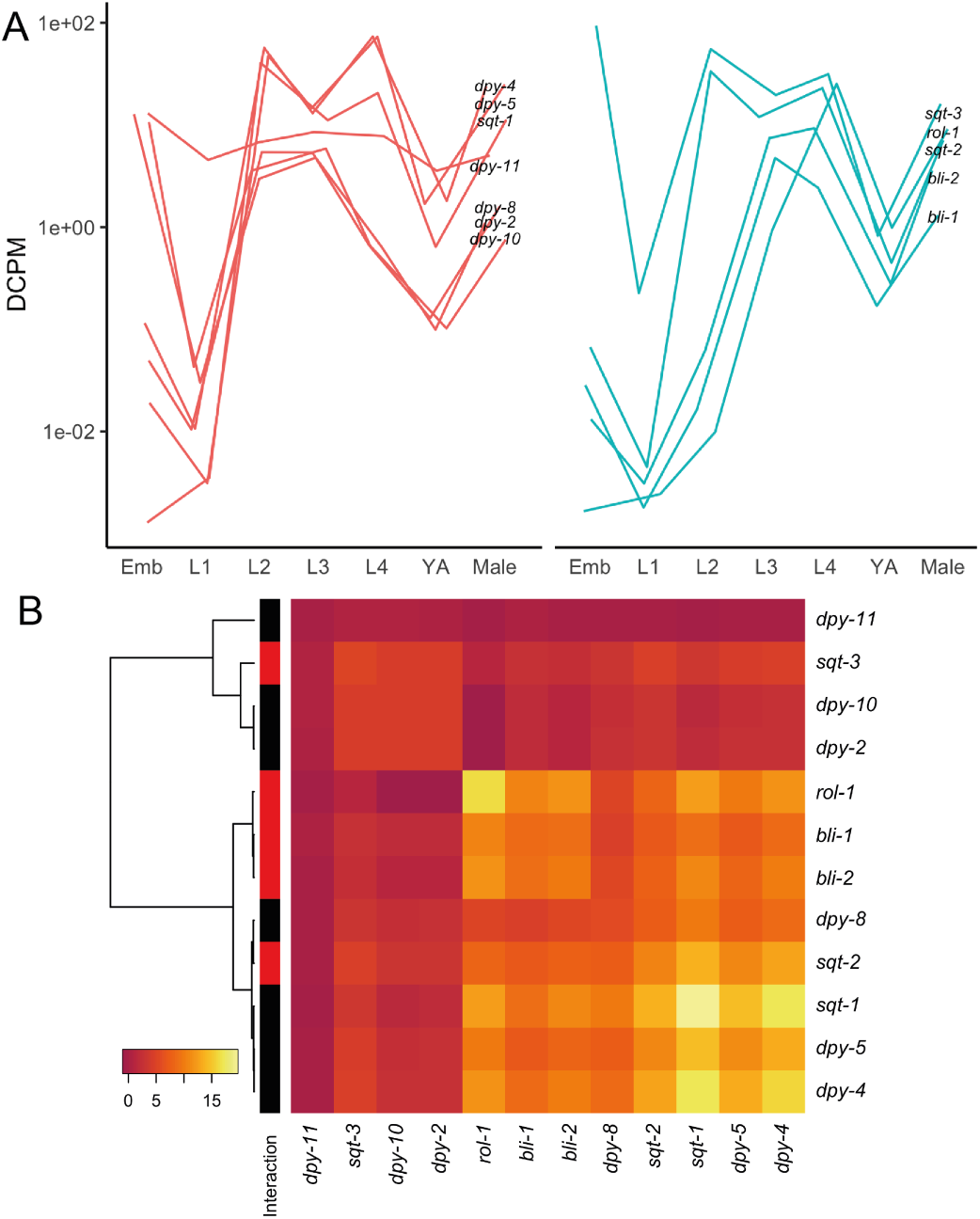
Developmental expression of cuticle genes, mutant alleles of which were tested for genetic interactions with *col-182*. Expression trajectories (**A**), split by the presence of any detected genetic interaction (*col-182* interactors at right, with a small positional jitter added along the x-axis as a visual aid), and expression covariance (**B**) across embryonic and larval stages and young adult (YA) hermaphrodites and males. Mean values of replicate experiments per stage are shown from Boeck *et al.* (2016).

*sqt-1, −2* and *-3* are all highly expressed collagens that interact genetically, with similar stage specificity from L2 onward (Figure 5). *sqt-3* is unique among collagens in its essentiality (Priess and Hirsh 1986; van der Keyl *et al.* 1994; Novelli *et al.* 2006), and is strongly expressed in the embryo as well. *sqt-1* also interacts genetically with *bli-1* and *bli-2*, and 22 other genes (Cox *et al.* 1980; Kusch and Edgar 1986; Kramer and Johnson 1993; Kramer 1994; Westlund *et al.* 1997; Nyström *et al.* 2002; Byrne *et al.* 2007; Shephard *et al.* 2011; Cai *et al.* 2011), yet we saw no obvious modification of the left rolling phenotype of *sqt-1(sc13)* mutants in the ancestral *col-182* background. This may be explained by the extreme specificity of allelic interactions among collagen mutants, and *sqt* mutants in particular. The *sqt-1(sc13)* allele tested is a recessive C-terminal C>Y substitution, altering cross-linking (Kramer and Johnson 1993; Yang and Kramer 1999), while *sqt-2(sc108)* is an N-terminal R>C substitution of unknown structural effect. Alleles of *sqt-1* vary markedly in their type, severity, temperature sensitivity, and degree of dominance of phenotypes, as well as inter- and intragenic interaction effects; from near wild-type for a null allele, to left or right rolling, abnormal hermaphrodite tail or male rays, or variation in body length. Enrichment in CeMEE interaction statistics for worm length/width over a small region spanning *sqt-1* (*p* < 0.00022) provides some fuel for speculation that allele-specific interactions with *col-182* may exist.

Lastly, no grossly visible interactions were seen for collagens involved in annuli formation and shape *dpy-2, −5, −8, −10* (McMahon *et al.* 2003). In sum, we surmise that *col-182* likely plays a role, apparently redundant under laboratory conditions, in one or both of the strut-anchored adult cuticular layers. The oft-touted genetic simplicity of *C. elegans* breaks down some-what when considering the cuticle, and targeted biochemical and structural analysis, together with epistasis analysis encompassing natural genetic variation, will be required to clarify the precise role of *col-182* and the majority of other collagens with no known function in the N2 background.

Effects of genetic background are ubiquitous in complex genetic systems wherever they are carefully considered. Studies mixing natural with domesticated genetic variation have amply shown the importance of genetic interaction on the phenotypic outcome of allelic effects in *C. elegans* (Seidel *et al.* 2008; McGrath *et al.* 2009; Bendesky *et al.* 2012; Duveau and Félix 2012; Gaertner *et al.* 2012; Andersen *et al.* 2014; Glater *et al.* 2014; Greene *et al.* 2016; Ben-David *et al.* 2017; Bernstein *et al.* 2018; Zhao *et al.* 2018). This extends to classical mutations of the cuticle, some of the first mutants isolated in *C. elegans* and core components of the worm geneticist’s toolkit.

## Acknowledgments

This work was supported by the National Science Foundation (DDIG 1210762 to TK), the National Institutes of Health (R01GM121828 to MVR), the NYU Dean’s Undergraduate Research Fund (AM), and the European Commission Sklodowska-Curie Fellowship (H2020-MSCA-IF-2017-798083 to LMN). For sharing data and facilities, we thank Henrique Teotónio. For strains, we thank Erik Andersen and the *Caenorhabditis* Genetics Center, funded by the NIH Office of Research Infrastructure Programs (P40 OD010440). We thank Jia Shen, John Yuen, Arielle Martel, James Hong, and Ambika Natesan for assistance with experiments.

## Notes

https://github.com/lukemn/cuticle

